# Iron chelation by curcumin suppress both curcumin-induced autoghagy and cell death together with iron overload neoplastic transformation

**DOI:** 10.1101/722942

**Authors:** Nathan E. Rainey, Aoula Moustapha, Ana Saric, Gael Nicolas, Franck Sureau, Patrice X. Petit

**Affiliations:** SPPIN – Saints Pères Paris Institute for the Neurosciences, UMR8003 CNRS, 45 rue des Saint-Pères, 75006 Paris, France; LVTS - Laboratory for Vascular Translational Science, UMR1148 INSERM, Université Paris XIII, Sorbonne Paris Cité, F-93017 Bobigny, France; Division of Molecular Medicine, Ruder Boškovic Institute, Bijenička cesta 54, 10000 Zagreb, Croatia; CRI - Centre de recherche sur l’inflammation, UMR1149/ERL8252 CNRS, Université Paris Diderot, faculté de médecine, 16 rue Henri Huchard, F-75018 Paris. France; LJP - Laboratoire Jean Perrin, UMR8237 CNRS, Université Pierre et Marie Curie, FRE 3231 Case Courrier 138, 4 place Jussieu, 75252 Paris cedex 05

**Author notes:** These authors contributed equally (Nathan E. Rainey, Aoula Moustapha). Correspondence &.

**Keywords:** Apoptosis, Autophagy, Cell death, Chelation, Curcumin, Endoplasmic reticulum, Ferroptosis, Iron, Mitochondria, Superoxide anions, Stress

## Abstract

Iron overload, notably caused by hereditary hemochromatosis, is an excess storage of iron in various organs which cause tissue damage and may promote tumorigenesis. To manage that disorder, free iron depletion can be induced by iron chelators like deferoxamine which are gaining interest also in the cancer field since iron stock could be a potent target for managing tumorigenesis. Curcumin, a well-known active substance extracted from the turmeric rhizome, has shown to be destabilizing endoplasmic reticulum and secondarily lysozomes, increasing mitophagy/autophagy and subsequent apoptosis. Recent findings show that cells treated with curcumin exhibit also a decrease in ferritin, which is consistent with it’s chemical structure and iron chelating activity. Here we investigated how curcumin would play on the intracellular effects of iron overload via Fe-Nitriloacetic acid or Ferric ammonium citrate loading in Huh-7 cells and explore consequences in terms of antioxidant activity, autophagy, or apoptotic signal transduction. With T51B and RL-34 epithelial cells experiments, we brought evidence that curcumin-iron complexation abolishes both curcumin-induced autophagy and apoptosis together with the tumorigenic action of iron overload.

## Introduction

Iron is a key element of numerous biological processes but the presence of free or loosely bound iron can be toxic to the cell (1). Iron being an active redox metal, the excess free form can generate reactive oxygen species through Haber-Weiss reduction followed by Fenton reaction (2, 3). Liver plays a major role in regulating iron storage in case of excess or deficiency in the blood pool. Aberrant iron accumulation can lead to cirrhosis and hepatocarcinoma but also heart damage, joint and metabolic disorders(4, 5). Phlebotomy remains the mainstay of iron-overload diseases but iron chelation therapy has proven to be a valuable alternative(6). Although iron chelation therapy was initially designed to alleviates toxic effect of excess iron occuring in iron-overload diseases, the novel toxicological properties of somes iron chelators complexes have radically shifted their intended applications toward cancer chemother apy (7–9). Some iron chelators are able not only to bind but also inhibit the redox properties of free labile iron. Such ligands may prevent iron from participating in Fenton reaction, inhibiting the formation of reactive oxygen species (ROS) like hydroxyl radical which initiates oxidative damages (10) and also ferroptosis, recently recognized as a form of regulated necrotic cell death (11). Ferroptosis is usually designed as one of the multiple variant of cell death than can be characterized by high intracellular level of free iron associated with ROS (12, 13). Ferroptosis is linked to the production of reduced glutathion and/or to the alterations of glutathion peroxidase 4 (GPX4) which usually acts as ROS controllers (14–16). Additives markers of ferroptosis are lipid peroxidation and protein carbonylation leading to more ROS and death signaling (10, 17). Interestingly, some genetic disorders of iron metabolism and/or chronic inflammation often evolve in iron overload and recent evidences show for example the possibility of lowering brain iron accumulation with the membrane permeant chelators deferiprone in Friedreich ataxia (18–21).

However, the choice of iron chelators is critical. Normally, the chelation of excess iron give rise to more inactive complex, hence providing a useful method to prevent toxic effect of iron-overload diseases. In contrast, some chelators are able to enhance the production of ROS after complexation with Fe^2+^. These chelators may provoke interesting Fe^2+^ toxicity for cancer chemotherapy (22, 23). By rapidly depleting proliferating cancer cells of Fe^2+^, iron chelators can also inhibit the activity of iron who acts as coenzymes to essential step of DNA synthesis (24–26). In addition, Fe^2+^ depletion are known to affect the expression of molecules involved in cell cycle progression and growth, i.e. N-myc downstream regulated gene 1, cyclin D1, cyclin A and p21waf1 leading to G1/S arrest in the cell cycle (27–29). These effects combined with ROS generation, provide multiple mechanisms of action mediated by the Fe^2+^ chelation to inhibit tumor growth.

Curcumin is the major chemical component of turmeric, a dietary spice made from the root of the Curcuma longa L. and used extensively in traditional Indian medicine (30). Curcumin is a potent bioactive compound actively in study against cancer (31–33), atherosclerosis(34), steatohepatitis (35) and neurodegenerative diseases,such as Alzheimer’s (36, 37) and Parkinson’s disease (38, 39), as well as to promote wound healing (40–42). At the cell level, curcumin mechanisms of action are complex and multifactorial. We and others previously address its mode of action and confirm its hormetic nature (43–47). Indeed, at low concentration, curcumin carries an effective antioxidant activity whereas at higher concentration (>20μM) behave as a potent pro-oxidant (47). In the field of cancer, curcumin inhibits the proliferation of tumor cells *in vitro* and *in vivo*. It inhibits also cell invasion, arrests cancer cells at G2/M phase of the cell cycle, and induces autophagy. Furthermore, curcumin suppresses the activation of Akt, mTOR and P70S6K proteins (46, 48, 49). Curcumin, therefore, is a potent tumor suppressor and simultaneously an inducer of autophagy. In case of failure, this sequence of events leads to apoptosis (44). All together, these data imply a fail-secure mechanism regulated by autophagy in the action of curcumin, suggesting a therapeutic potential for curcumin offering a novel strategy for the treatment of malignant cells (47). Curcumin modulates proteins of iron metabolism in cells and in tissues, and chemical structure suggests that curcumin has properties of an iron chelator (50–53). Indeed, the *α*,*β* unsaturated diketone moiety of curcumin can form chelates with transition metals (Figure 1) and especially with iron (51, 54)[51]. Metal chelates of curcumin are mostly non-fluorescent, although absorption spectra shows significant changes that can be tracked for assessing efficient chelation. Metal chelates of curcumin of the type 1:1 and 1:2 have been reported for the ions Cu^2+^, Fe^2+^, Mn^2+^, Pb^2+^, and others (55). All these metals can play a role in amyloid aggregation, and curcumin is investigated as a chelating agent to act on Alzheimer pathogenesis (54). Since curcumin is also lipophilic and readily crosses membranes it may also chelate more metal ions intracellularly(56). Evidence shows that curcumin is more efficient when low amount of iron are present intracellularly (53). However, how curcumin binds to these metals was not investigated in details and neither the consequences of this interaction in terms of autophagy and cell death mechanisms.

**Fig. 1.**
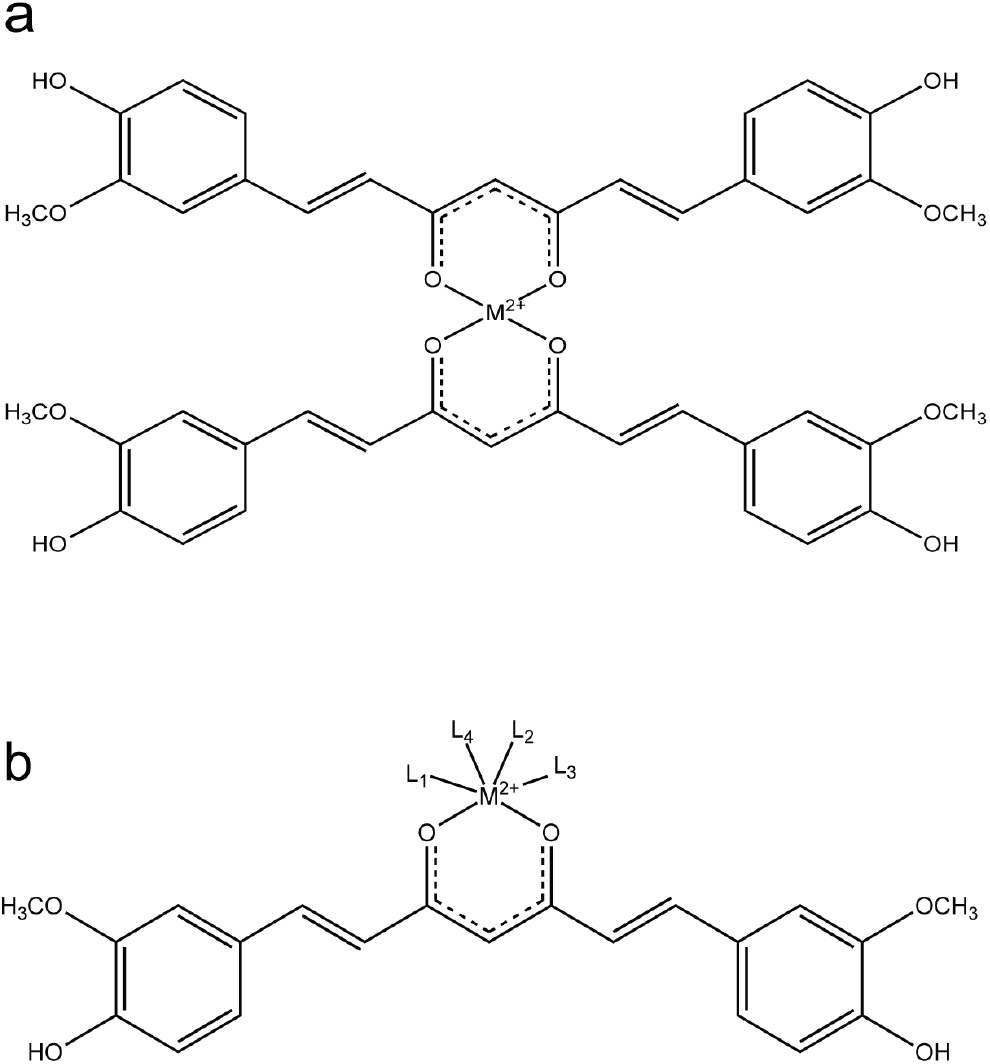
Curcumin chelates metals. Schematic representation of the curcumin chelation towards bivalent metals | **a.** Curcumin chelates can adopt a 2:1 configuration | **b.** Curcumin can also chelates metals in conjunction with other ligands

In this context, curcumin may be effective in preventing chemically-induced liver damages, by decreasing liver toxicity in rats in the case of galatosamine or carbon tetrachloride intoxication (57) and also abolished carcinogenic effects of afflatoxin and/or nitrosodiethyamine (58, 59).

Here we demonstrate that curcumin do not change the amount of intracellular iron loaded but that it’s chelating activity is crucial for inhibiting iron overload cytoxicity and further effects on liver cell line. These findings put curcumin as a powerful alternative candidate for the control of liver damages linked to intracellular iron overload. Moreover, we tested the capability of curcumin as powerful iron chelator to prevent tumor promotion in T51B and RL-34 cells. Together with the fact that curcumin binds to iron and prevent iron toxicity (60–62), we find that curcumin chelation act similarly as deferroxamine (29) by inhibiting tumor promotion.

## Results

### Curcumin binds iron

Evidence for iron binding to curcumin in solution has been previously demonstrated using spectroscopic shift techniques (54, 63). The Figure 1 shows capability of curcumin to chelate divalents cations like Fe^2+^ directly (1a) or in conjunction with other ligands (1b). We used then, an alternative metal affinity chromatography method with handmade Fe-NTA columns. The Figure 2a shows the loading efficiency of curcumin from solution to the Fe-NTA-agarose column. Curcumin binding was rapid and efficient, reaching up to 90-95% after incubation for 10 minutes on the iron-containing resin. It is well known that the diketone groups of curcumin likely interact with the Fe-NTA-resin in a similar fashion than the phosphate oxygens of the phospho-peptides, for which this iron affinity chromatography resin was developed (64). The data extracted from Figure 2 gave an affinity estimation around μM, in agreement with the high affinity iron binding site of curcumin reported by Baum et al. (54). Comparable curcumin binding was not observed when Fe was substituted by Ni as the metal ion (64). A significant fraction of curcumin (based on absorbance signal at 435 nm) was recovered from Fe-NTA-agarose upon addition of competitive iron chelators like deferoxamine (DFO) or EDTA (EGTA to a lesser extent)(Figure 2b). Vitamine E has much less efficiency of being an iron chelator. These data reinforced previous reports of curcumin specifically binding iron, and suggests may suggest a potential to reduce both uptake and toxicity of iron within cells. The curcumin absorbance spectra is modified by the presence of Fe-NTA. Effectively, the absorbance of curcumin shifted from 435 nm to 400 nm for the complex when bound to iron (Figure 2c). Natural fluorescence behavior of free curcumin peaking at 535nm is completely quenched when incubated with Fe-NTA, either in emission spectra or microspectrofluorimetry on cells (Figure 2d,e). Acellular experiments also demonstrate that Fe-NTA added to curcumin provoke an instantaneous, dose dependent quenching of curcumin fluorescence (Figure 2f).

**Fig. 2.**
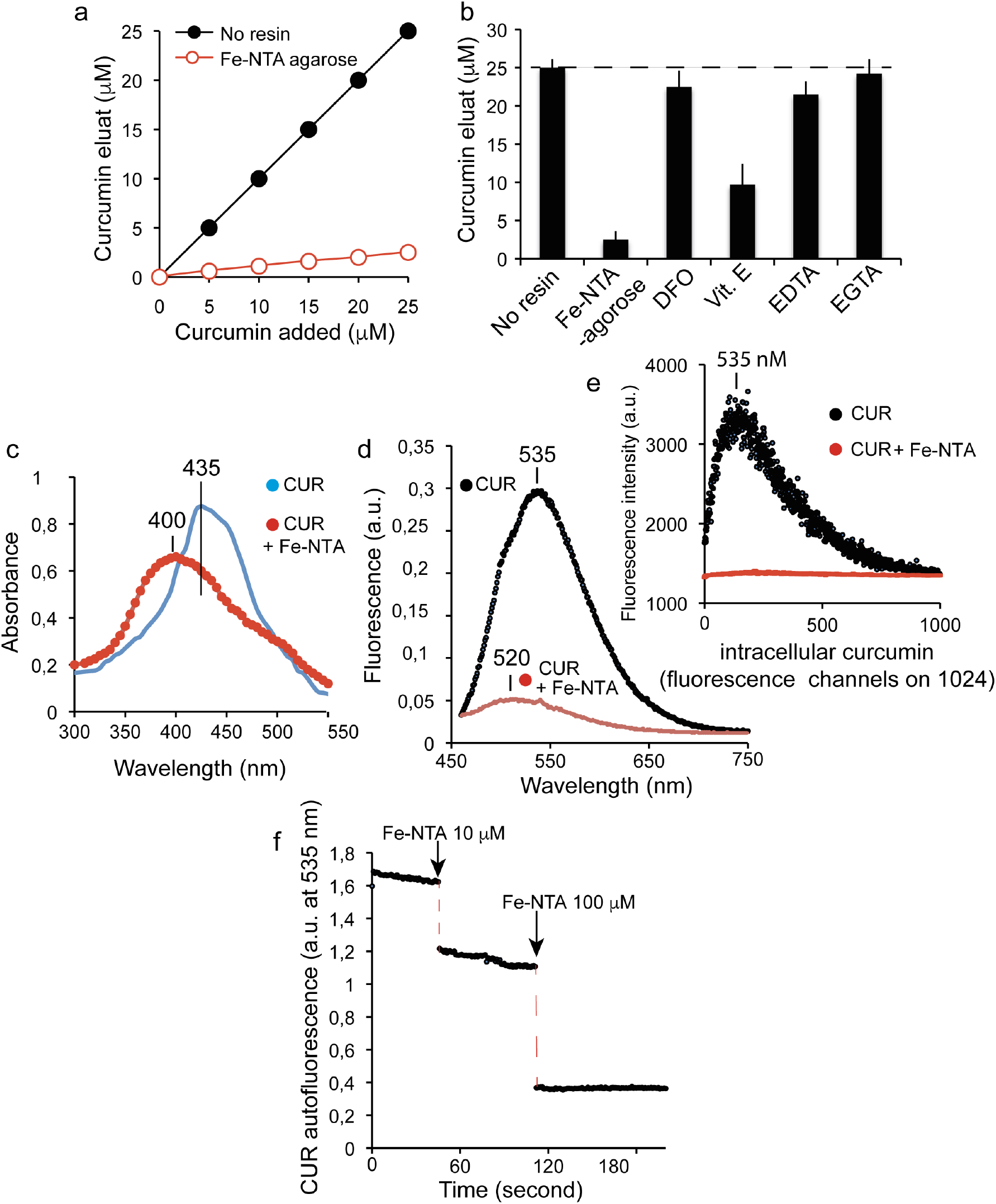
Curcumin binds iron. **a.** Test of the removal of curcumin from solution by Fe-NTA resin. Increasing amounts of curcumin were incubated with Fe-NTA-Agarose (black circles) or without (open circles). The free curcumin remaining in solution after removal of the resin was determined by absorbance at 435 nm against calibration range | **b.** Inhibition of curcumin binding (20μM) incubated alone (no NTA resin) with iron resin (Fe-NTA agarose) or with iron resin pre-incubated with excess deferoxamine (DFO 250 μM), excess vitamin E (Vit.E 1 mM), excess EGTA (2 mM) or excess EDTA (2 mM). Unbound curcumin was determined as for panel a. Means ± SE are presented from at least five independent determinations | **c.** Absorbance spectrum of Huh-7 cells treated by of 20 μM curcumin for 6h with (in red) or without (in blue) 100 μM Fe-NTA loading for 24h | **d.** Fluorescence emission spectra of Huh-7 cells treated by of 20 μM curcumin for 6h (same conditions as panel c)| **e.** Microspectrofluorimetry of Huh-7 cells treated as panel (c,d) and recording of the curcumin emission | **f.** Effect of Fe-NTA addition on the fluorescence recording of a curcumin solution (20 μM) at 535 nm realized in cuvette with an excitation at 435 nm.

### Curcumin efficiently chelate Fe^2+^ but does not block iron uptake in Huh-7 cells

The dosage of intracellular iron shows that iron chelation by curcumin do not alter cellular iron uptake whereas curcumin only slightly affect the total amount of intracellular iron (Figure 3a,b). The detected cellular iron is in the range of 28 ± 10 μg/g of cells for a 48h incubation with 100 μM Fe-NTA whereas control cells contained 7,2 ± 1,9 μg iron/g of cells (Figure 3b). A pre-incubation with DFO and CoCl_2_(data not shown) almost totally abolish iron content of the cells treated with 100μM Fe-NTA (Figure 3b). Meaning that DFO and CoCl2 act mostly on the iron transport activity when curcumin acts intracellularly with an almost strict chelating activity (immobilization of the free intracellularly available Fe^2+^). Additive experiment with Calcein, whose fluorescence quenching is strictly correlated with intracellular accumulation of iron, first demonstrates efficient quenching by Fe-NTA treatment and FAC (Figure 3c). Also, whatever the treatment method used, curcumin complexation with iron restore calcein fluorescence (Figure 3c).

**Fig. 3.**
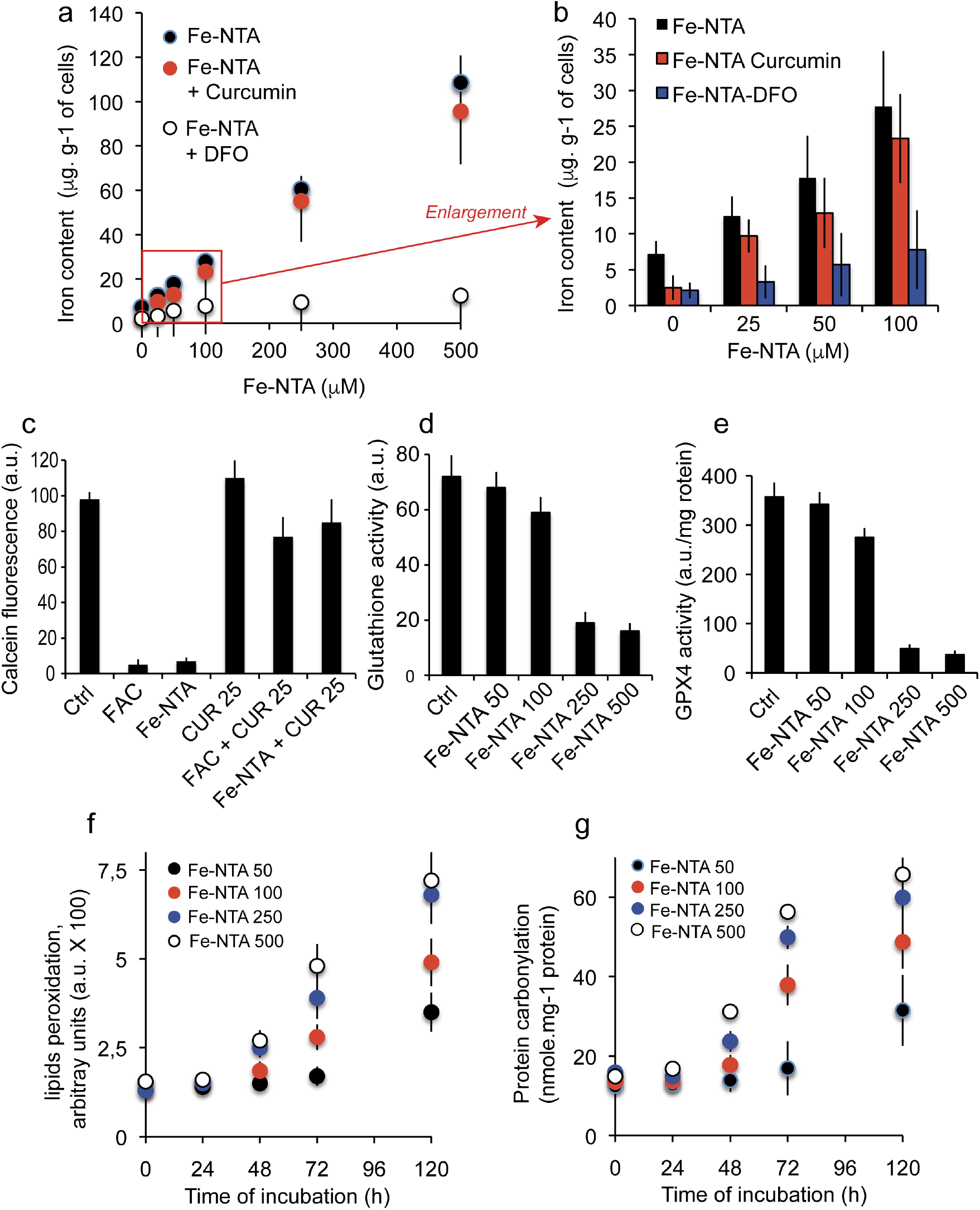
Cellular iron content & oxidative consequences. **a.** Fe-NTA cell loading (0-500 μM) and measurements of the intracellular iron content in presence of DFO or Curcumin (20 μM) | **b.** Enlargement of the Fe-NTA cell loading in the lower range between 0 and 100 μM in absence or presence of DFO (excess DFO, 250 μM) or curcumin (20 μM) | **c.** Cells were charged with calcein-AM and either FAC (200 μM) or Fe-NTA (100 μM) then compared at cells loaded with curcumin only (20 μM). On last conditions cells are simultaneously treated with FAC and curcumin or Fe-NTA and curcumin. After 1h the cells are rinsed with phosphate buffer saline (PBS) on ice and the calcein fluorescence measured (± SD for 6 repeated experiments) | **d.** Effect of various treatment of Fe-NTA (0 to 500μM) on the glutathion activity of Huh7 cells | **e.** Same experiments as panel d. for assessing Glutathion peroxidase (GPX4) activity| **f.** Effect of iron loading (as panel d) on lipid peroxidation as a function of time (0-120h) | **g.** Effect of iron loading (as panel d) on protein carbonylation as a function of time (0-120h).

### Toxicity features associated to Fe-NTA loading

Increased loading of Fe-NTA (from 50 to 500 μM) is linearly related to the intracellular iron content but also associated (at doses ≥ 100μM) with a significant reduction of the glutathion content (Figure 3d) and glutathion peroxidase activity (Figure 3e). These events are associated with lipid peroxidation and protein carbonylation (Figure 3f,g). However, at the concentration of 100 μM Fe-NTA for 48h that we currently used in our work all these four parameters are discrete.

### Cell death induction by curcumin

Huh-7 cells treated with curcumin at 25 μM for 48h and double-stained with YO-PRO-1 versus PI exhibited different populations including viables cells (Y−/PI−), apoptotic cells (Y+/PI− or Y+/PI ±, i.e. slightly positives) and cells undergoing necrosis (Y+/PI+++). The maximal apoptosis being reached at 25 μM for 24h since higher curcumin concentrations favor necrosis (Figure 4a). It is interesting to notice that there is a huge difference betwwen 10 μM and 25 μM curcumin treatment and also that cumulated cell death (apoptosis + necrosis) reach 60 to 70% of death at 25 μM for 24h. Clearly that is not the case with curcumin 10 μM where the overall dead cells only reached 7% at 48h ((Figure 4b, data not shown). It is not surprising, since curcumin binds to the ER, that the apoptotic processes are linked to calcium release from the endoplasmic reticulum (Figure 4c) during the first 4 hours of curcumin internalization. This calcium release influence directly the mitochondrial behavior, so, the mitochondrial membrane potential drop together with an increase production of superoxide anions probably associated with the mitochondrial permeability transition pore (MPTP) opening(Figure 4c). Following that ΔΨ*m* drop, an important caspase-8 and caspase 3/7 activation occur as well as an NAD(P)H and NADH (fluorescent form) switch towards the reduced forms, NAD+ and NADP+ (which are not fluorescent), then resulting in a drop of NAD(P)H that paralleled the ΔΨ*m* decrease of fluorescence (Figure 5c). Clearly, if a comparison is made between the curves of apoptosis/necrosis and the physiological events underneath, their is a correlation between the disruption of the mitochondrial homeostasis (i.e. ΔΨ*m* drop and O_2_.-production) and the early increase of apoptotic events followed by secondary necrosis (Figure 4b). The curcumin loading is quite rapid and associated with an immediate increase in red acridine orange fluorescence indicating an early induction of autophagic processes (Figure 4d). The correlation between AO red fluorescence and LC3-II occurrence has been previously described [44] and suggest that at certain concentrations, primary events induced by curcumin are linked to autophagy.

**Fig. 4.**
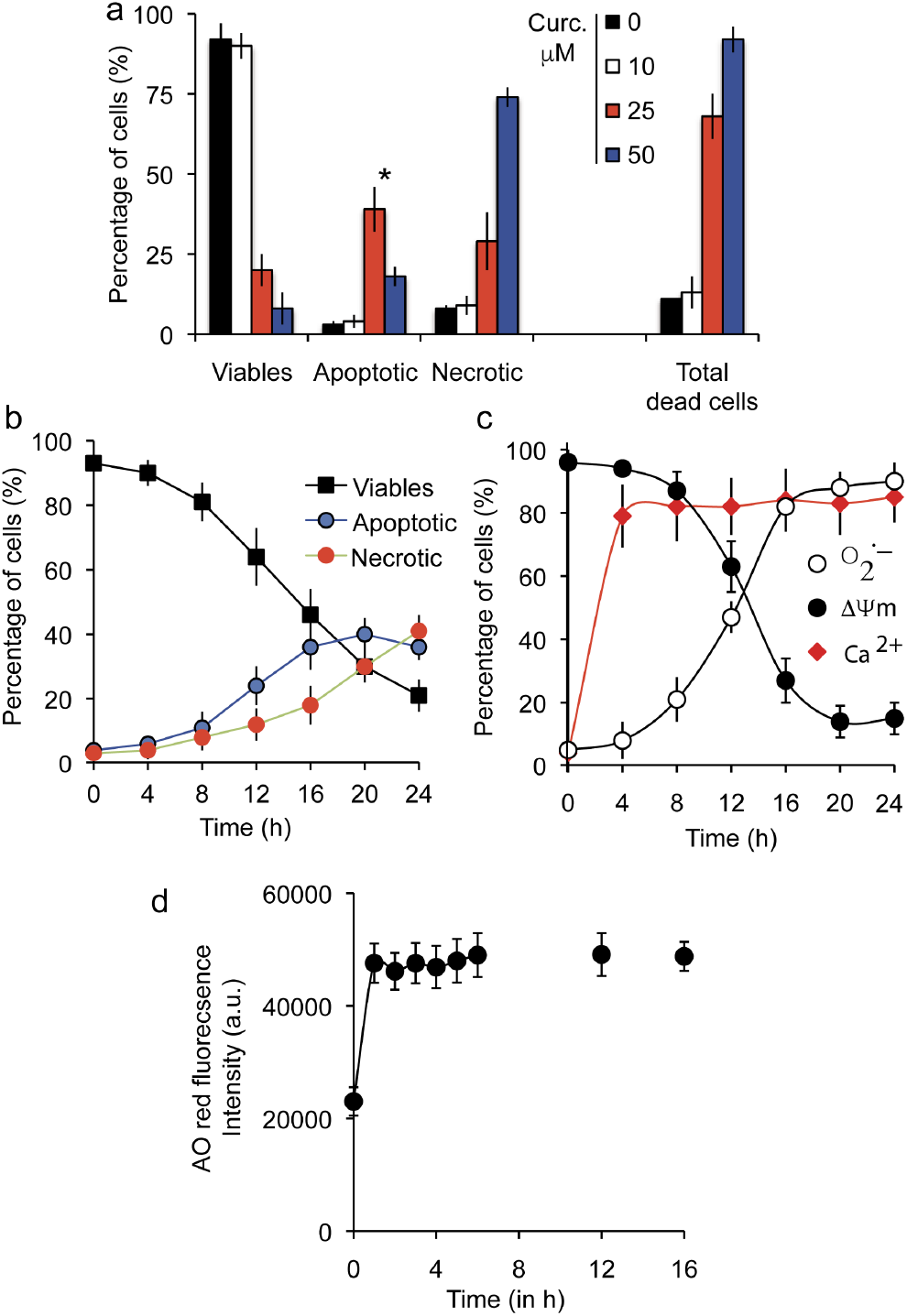
Prototypic induction of cell death by curcumin. **a.** Flow cytometry analysis of Huh7 cells treated with various concentrations of curcumin (0-50 μM for 24 h incubation). YOPRO-1 / PI staining was used to analyze membrane permeability and discriminate three populations: viable (YOPRO-1− / PI−), apoptotic (YOPRO-1+ / PI− & YO-PRO-1+/PI±) and necrotic cells (YOPRO-1+ / PI+) **\*** The maximal amount of apoptotic cells is observed at 25 μM of curcumin for 24 h. Data are expressed as the mean ± S.D. (n=7) | **b.** Same experiment as panel a but plotted as a function of time from 0 to 24h (curcumin 25 μM) | **c.** Metabolic measurements that are linked to the dying cells after 24 hours incubation at 25μM curcumin. Mitochondrial membrane potential with DiOC_6_(3) fluorescence (ΔΨ*m*, black circles), superoxide anion generation tested with MitoSOX-green fluorescence (O_2_^.-^, empty circles) and calcium rise measured with Fluo 4-AM (intracellular calcium, red diamonds) | **d.** Acridine orange (AO) staining of cells during the 16 first hours after 25 μM curcumin. Increased staining with rise of low pH vesicles (lysosomes) and fusion with bigger vesicles (autophagosomes) can be associated to autophagy induction.

**Fig. 5.**
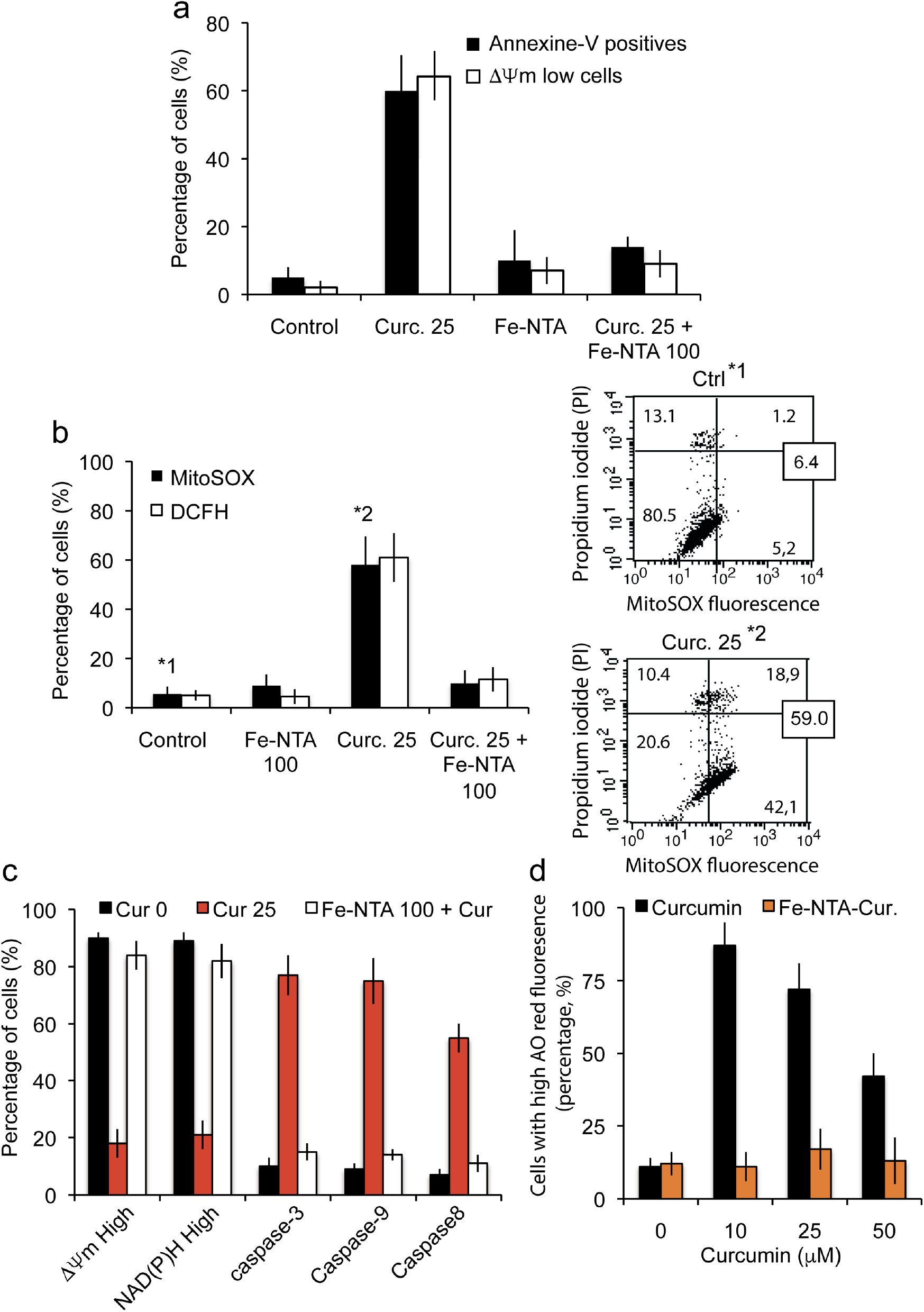
Fe-NTA-curcumin complex abolish their own toxicity. **a.** Flow cytometry analysis of early and later events towards cell death (respectively drop of ΔΨ*m* and PS exposure) when cells are treated for 24 h with 20μM curcumin or 100μM Fe-NTA alone or both products behaving as a chelate | **b.** Flow cytometry analysis of the consecutive events to the ΔΨ*m* drop (panel a) that are the generation of superoxide anions (MitoSOX staining) and hydrogen peroxide production (DCFH-DA staining; each histogram is taken in 7 independent experiments and ± SD is given). Control and 25μM curcumin (24h) conditions are illustrated respectively in 1* and 2* by biparametric plot together with the viability measurement (PI staining). The percentage of cells in each panel is given as a percentage of the whole population | **c.** Metabolic analysis showing that treatment with both products restore control condition trend with respectively a large population of high ΔΨ*m* (baseline) and NAD(P)H together with a low activation of initiation caspase (caspase-8) and executioner caspases (caspase-3 and caspase-9). Cells were treated with 25μM curcumin or 100μM Fe-NTA or both | **d.** Cells were treated for 24h with curcumin or Fe-NTA, or both. Acridine orange (AO) staining is performed to investigate AO red fluorescence related to vesicle fluxes. Fe-NTA-curcumin complex abolish the increase caused by curcumin alone and run at baseline level.

### Iron chelation with curcumin resulted in a full inhibition of curcumin-induced autophagy and apoptosis

As demonstrated above, the cells treated with curcumin alone are undergoing autophagy (Figure 4d), apoptosis and necrotic death secondary to apoptosis (Figure 4a,b) with is characterized by a progressive loss of viability, a drop in ΔΨ*m* associated with an increase of superoxide anions production but also the initiation of caspase-8 together with caspase-3 and caspase-9 (Figure 5c). Whereas, the iron-chelated with curcumin abolished completely the cell death inducing capacity of curcumin. Indeed, the expression of phosphatidyl serine at the cell membrane (late apoptotic and necrotic event) is abolished as well as the ΔΨ*m* drop usually detected as an early event (Figure 5a). Superanions and hydrogen peroxide production are abolished (Figure 5b). Also, NAD(P)H fluorescence and caspase-3/7, −8 and −9 are not activated anymore as curcumin alone do (Figure 5c). Moreover, the AO red fluorescence linked to the increasing amount and big ger size of acidic vesicle linked to autophagy enhancement are maintained to control level (Figure 5d), whatever the curcumin concentration and up to 50 μM. Curcumin-induced autophagy is decreasing with increasing curcumin concentration because apoptosis and necrosis becoming predominant for treated cells (Figure 5d). It could be observed that below 10 μM, curcumin usually do not promote cell death but that autophagy is engaged. Above 10 μM, the amount and intensity of AO positives cells is decreasing as apoptosis and secondary necrosis take place.

### Chelating activitity of curcumin on iron alleviates its tumor-promoting effect

To assess if curcumin, as a potent iron chelator, can also reduce or prevent the tumor-promoting effect of iron overload, we used T51B (rat liver epithelial cell) and RL-34 (rat liver epithelial diploid cell line). We used an impedancemetry control system (Xcelligence, ACEA) to follow and synchronize pre-culture before seeding on soft agar,and have decided to take the cells after 36h when they are in the maximal proliferation phase (Figure 6c, black arrow). This has been done for both T51B and RL-34 cell lines. Previous studies show that iron act as a tumor promotor in T51B cells with a FAC induction associated to a low dose of N-Methyl-N’-nitro-N-Nitrosoguanidine (MNNG) generally acting by successives cycles of hepatotoxicity followed by regrowth and regeneration [71]. Acute iron overload is done with 250μM Fe-NTA or with FAC + 8HQ for 7 days in absence or presence of increasing amount of curcumin. The two cell lines exhibit a similar behavior and 50μM of curcumin is enough to efficiently counteract iron overload toxicity (Figure 6a,b). Whatever the iron overload protocol is, the induction of colonies takes over 12 weeks of repeated cycle of cells culture on soft agar, and the number of colonies is quite significant at 20 weeks of culture (over 120 colonies of 25.000 cells) (Figure 6d). By chelating iron, curcumin can control the number of colonies development who do not go over 5 colonies after 20 weeks of culture (Figure 6d) in both cell lines.

**Fig. 6.**
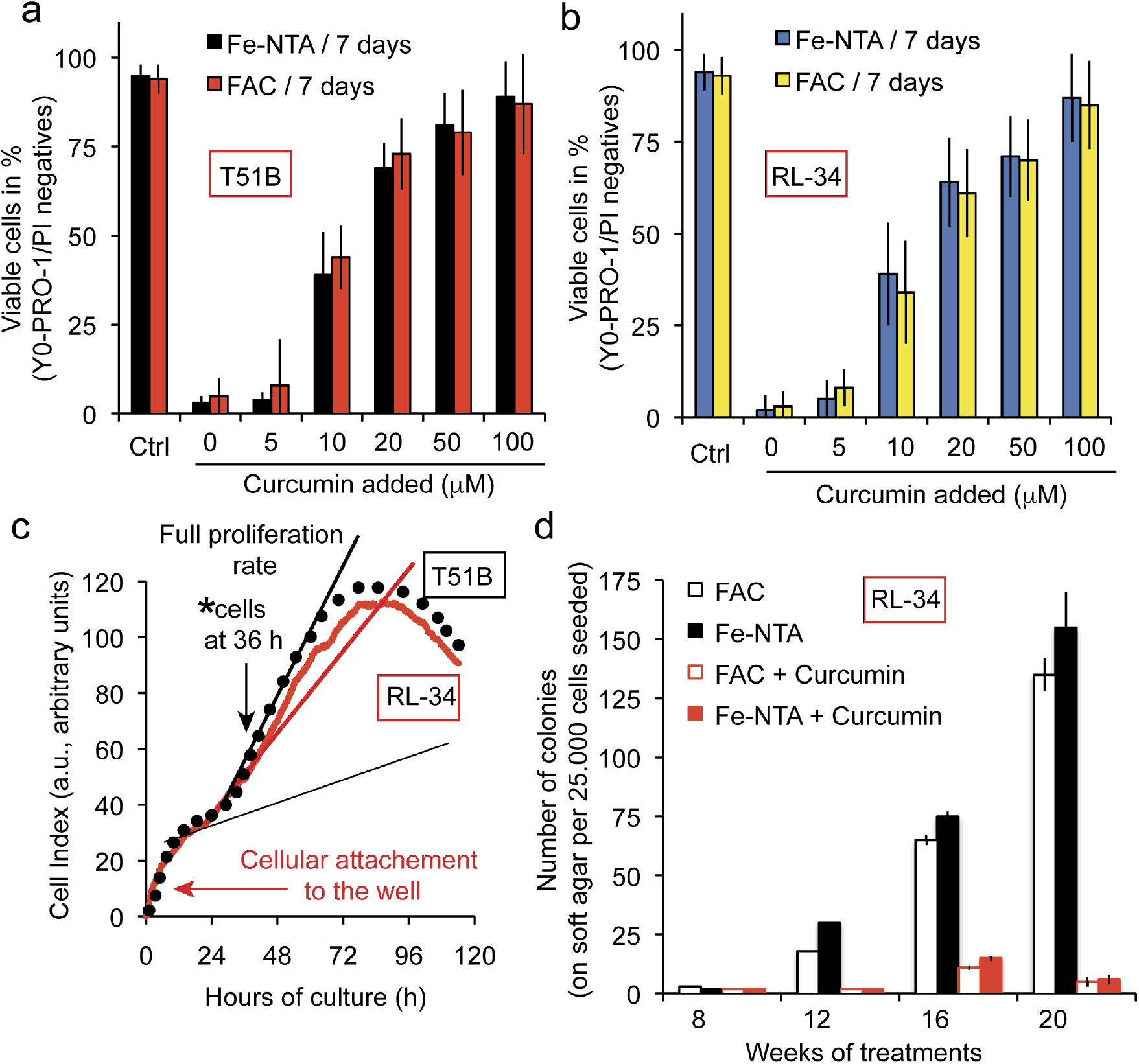
Curcumin chelate with free iron alleviate the tumor promoting activity Fe-NTA or FAC. **a.** T51B cells are protected against free iron toxicity by curcumin in a concentration-dependant manner. Epithelial T51B cells were treated 7 days with either Fe-NTA 100μM or FAC 100μM (+10μM hydroxyquinoline) and various curcumin concentrations and then viability-tested by flow cytometry taking into account the dead cells (apoptotic, A and necrotic, N cells) to find the cellular viability (100 - (A+N) = Viable cells in %) | **b.** Same as in panel a but with RL-34 cells | **c.** The cells used in panel d experiment are first followed by impedencemetry (ACEA Xcelligence system) and taken at 36h in their full proliferation phase after plating. Arrows (black) are setting the situation of the cells at 36 h and the Arrow (in red) along the early time on the curve pointed out the initial cell binding to the wells that in our case take almost 24h before full proliferation take place. The curve is the mean value of 12 differents wells | **d.** T51B cells plated at 2.10^5^ cells in 60mm dish were treated with 0.5 mg/ml MNNG for 24 hours after plating and cultured either with 20 μM curcumin or Fe-NTA 200 μM (or FAC 250 μM with 8-hydroxyquinoline), or both. The number of soft agar experiments conducted was n= 7 and ± SD were taken.

## Discussions

### Curcumin to iron interactions

Previous experiments by cyclic voltammetry showed that curcumin binds metal ions and especially Fe^2+^ (63). The iron chelation activity of curcumin is related to it’s chemical structure via the *β*-diketone group, a known bidentate chelator of Fe^2+^, similar to group found in simple analogous complex of Fe(II) and acetylacetonate. Based on the formation constant of Fe(II)-curcumin of 10^22^ M^−1^, it has been calculated that the pFe^2+^ (pM) of curcumin is of 16.6 (at pH 7.4 for 10 μM curcumin and 1 μM Fe^2+^)(63, 65). This curcumin pM is not as high as other iron chelators already used in iron-overload treatment such as deferiprone (pM = 20) and deferoxamine (pM = 26) [73]. Curcumin pM could be compare favorably to the pM of the iron chelator nitrilotriacetic acid (NTA) that we use on the nickel column for curcumin extraction and many other iron chelators (Figure 2) and fit well with other iron chelating activity detected *in vitro* (65).

The methods used for loading iron to the cells allows us to go from an endogenous iron content of 7,2 ± 1,9 μg iron/g of cells to a content of 28 ± 10 μg/g of cells for a 48h incubation with 100 μM Fe-NTA (Figure3a). These results are not hindered by deleterious effects of iron as toxicity features become significative only above 100μM and longer time period. Iron toxicity is featured with lipid peroxidation and protein carbonylation (Figure 3 f,g) also associated to a decrease in intracellular protection against ROS like glutathion activity decrease (Figure 3 d,e). The iron accumulation at higher Fe-NTA concentrations for 48h leads to an intracellular accumulation of 60.5 ± 13.2 and 108.5 ± 19.6 μg/g for Fe-NTA 250 μM and 500 μM, respectively. These conditions of loading go beyond the capacity of the endogenous Antioxidant defense system (ADS) with a clear depletion of the glutathion activity, overwhelmed by massive generation of ROS from cellular Fe^2+^ engaged in Fenton reaction / Haber-Weiss effect (2, 3). Despite curcumin clearly bond with iron, there is no significant reduction of the iron uptake within cells (Figure 3a) although we could consider a discrepancy due to transient interaction between curcumin and plasma membrane during primary penetration where it could interact with iron regulation system. So, from our data it results that curcumin binding to iron do not influence the intracellular iron amount in response to Fe-NTA treatment. We then, investigate if curcumin chelation with iron still feature an hormetic behavior or if it’s activity is altered.

### Curcumin cellular effects

We investigated the Huh7 profile response to curcumin as we previously done (44) and confirmed that curcumin exert an effect on endoplasmic reticulum which release calcium taken up by mitochondria via mitochondrial calcium uniport (MCU) who cause a drop in ΔΨ*m* and a superoxide anions production, leading ultimately to an opening of the mitochondrial permeability transition pore (mPTP) (Figure 4c). These results fit with the further involvement of cathepsins and caspase-8 activation leading to cytochrome c release and apoptosis previously described (44). This do not exclude also en early involvement of lysosomes as it is clear that curcumin in the μM range induce autophagic processes (Figure 4d)(44). This may explain some observations made by colleagues about the acceleration of neoplastic transformation when curcumin in the range of 10-20 μM is added to iron (29, 66). It is possible that 10-20μM of curcumin is more than enough to chelate all iron, and the remaining iron-free curcumin induce autophagy, enhancing neoplastic transformation (44, 47). Indeed, in neo-plasic cells, autophagy constitute a coping mechanism with intracellular and environmental stress, thus favoring tumor progression (67).

### Curcumin-iron chelates abolish both curcumin effects and iron cytotoxicity

When chelated with iron, curcumin loose capability to induce toxicity and cell death (Figure 5) which is in line with previous reports showing that iron attenuate curcumin cytotoxic effects in squamous cell carcinoma (68). That chelation is able to abolish late stage signals like outer exposure of phosphatidyl serine (Figure 5a) but also early production of both hydrogen peroxides and superoxide anions from the mitochondrial compartment (Figure 5b). These results fit w ith o ther i nhibition l inked t o c ell d eath l ike the drop of mitochondrial potential, reduction of NAD(P)H, activation of caspases 3/9/8 (Figure 5c). These results contrast with the ongoing hypothesis that combination of ions and ligands would produce a synergistic effect on cells (69–71). A possible explanation would come from the fact that the central carbon has a labile hydrogen that is locked into the chelate and is truly unable to produce oxyradicals unless complete dissociation. According to these data, keto-enol function is more likely to drive the curcumin-induced autophagic process (Figure 5d). Another hypothesis to consider would be the modification of the global size and hindrance of the curcumin chelate compare to free curcumin.

Soft agar tumorigenesis model allowed us to investigate for a longer period how RL-34 cells would evolve on iron-induced stress (with a MNNG starter) and curcumin treatment. As FAC or Fe-NTA treatment (250μM) are able to produce large number of colonies, 20μM curcumin is enough to abolish that growth (Figure 6d). Messner et al. recently evidenced this same response for 20μM curcumin but observe a tumor promotion for 10μM curcumin in the same settings (66). We hypothesize that curcumin at 10μM is already in excess compared to free iron, especially if curcumin can cycle the iron to transferrin and maintain it in redox inactive state (52). That excess is not enough to induce apoptosis and control tumorigenesis like 20μM does (Figure 6d), but enough to stimulate autophagic processes, hence, boosting tumorigenesis. On clinical cancer settings, we would need to address a specific curcumin payload, either partially chelating iron to reduce this tumor-promoting signal or increase payload enough to produce apoptosis and not boosting autophagy. On the opposite, iron-overload diseases management don’t suffer that limitation, and while iron chelation is seeked, an extra autophagic signal could be valuable for patients at the tissue level, as well as anti-oxidative properties provided also by low concentration of curcumin.

## Methods

### Chemicals and reagents

Curcumin, deferoxamine, 8-hydroxyquinoline (8-HQ), ferric amonnium citrate (FAC), propidium iodide (PI), N-acetylcysteine (NAC) were obtained from Sigma-Aldrich Chemical Co. (St. Louis, MO, USA). Calcein-acetoxymethyl(Calcein-AM), 2,7-Dichlorodihydrofluorescein diacetate (DCFH-DA),3,3’-dihexyloxacarbocyanine iodide [DiOC_6_(3)] and N-[4-[6-[(acetyloxy)methoxy]-2,7-difluoro-3-oxo-3H-xanthen-9-yl]-2-[2-[2-[bis[2-[(acetyloxy)methoxy]-2-oxoethyl]amino]-5-methylphenoxy]ethoxy]phenyl]-N-[2-[(acetyloxy)methoxy]-2-oxoethyl]-,(acetyloxy)methyl ester (Fluo-4 AM) were purchased from Molecular Probes (Invitrogen, Eugene, OR, USA). Agarose was obtained from Lonza (Walkersville, MD, USA. The 8-hydroxyquinoline is given together with FAC to enhance its internalization.

### Cells

Human hepatoma-derived Huh-7 cells (RIKEN BioResource Center, Tsukuba, Japan) were grown in the presence of 5% CO2 with Dulbecco’s modified Eagle’s medium (DMEM) containing high glucose (25 mM Sigma-Aldrich, St. Louis, MI) with 10% Fetal Bovine Serum (FBS, Hyclone, Logan, UT) completed with 1% penicillin-streptomycin, HEPES NaOH 1 mM, Na-pyruvate 1 mM and 1% nonessential amino acids (MEAM, GIBCO). T51B cells (rat epithelial cells lines, non-neoplasic) are a model to study tumor promotion in vitro. They can be transformed to grow in soft agar by treatment with low amount of carcinogens and tumors promotors (72). RL-34 cells (rat liver epithelial-like cells, non-neoplastic from JCRB bank, japan) were maintained with the same medium than T51B cells.

### Neoplastic transformation

T51B and RL-34 cell neoplastic transformation was realized as described by Messner et al.(66) and the formed colony on soft agar counted as a transformation index(72). We follow exactly previous transformation protocol with a single treatment of 24h with 0.5 μg/ml N-methyl-N’-nitrosoguanidine (MNNG) followed by continuous culture for 20 weeks either with FAC (+ 8-HQ) or Fe-NTA at 250 μM with or without curcumin at 25 μM. MNNG is added at the start to participate to the initiation of the tumorigenesis. The culture media is renewed almost every 2-3 days and colony formation are tested every two weeks with a special care when becoming more important at 16, 18 and 20 weeks of culture.

### Intracellular Iron dosage

Intracellular iron content of Huh7 cells given in micrograms of iron per grams of cells were determined after treatment with 100μM Fe-NTA or/and 100μM DFO for 24h. 150cm^2^ cell culture were trypsinized and collected with lysate buffer (trichloroacetic acid 0,1g/L, chlorhydric acid 0.774M, Mercaptoacetic acid 30mM) to respect a 100mg cells for 400μL buffer. Lysate are heated at 65°C overnight and centrifuged. Supernatants are then analyzed by flame emission spectrometry (Institut Claude Bernard, ICB, Paris) and compared to standards to determine iron content.

### Microspectrofluorimetry

The UV-visible confocal laser microspectrofluorimeter prototype was built around a Zeiss UMSP80 UV epifluorescence microscope (Carl Zeiss, Inc., Oberkochen, Germany), optically coupled by UV reflecting mirrors to a Jobin-Yvon HR640 spectrograph (ISA, Longjumeau, France). The 351nm UV line of an argon laser (model 2025; Spectra-Physics, Mountain View, CA) was used for either drug or fluorochrome excitation. The diameter of the laser beam is first enhanced through a double-lens beam expander in order to cover the entire numerical aperture of the microscope’s optics. The laser beam is then deflected by the epi-illumination system (dichroic mirror or semireflecting glass) and focused on the sample through the microscope objective (X63 Zeiss Neofluar water-immersion objective; numerical aperture = 1.2) on a circular spot 0.5 μm in diameter. The excitation power is reduced to less than 0.1 mW by neutral optical density filters. The objective was immersed in the culture medium, and a circular area 0.8 μm in diameter was selected at the sample level, by interposing a field diaphragm on the emission pathway of the microscope, to selectively collect the fluorescence signal from the nucleus or a specific cytoplasmic area. Confocal conditions are met when the image of this analysis field diaphragm through the microscope objective perfectly coincides with the focus of the laser beam on the sample. Under these conditions, the experimental spatial resolution, measured on calibrated latex beads (2, 0.6, and 0.16 μm in diameter) labeled with the fluorescent probe fluorescein, is 0.5 μm for the directions X, Y, and Z. Finally the fluorescence spectra were recorded after spectrograph dispersion, in the 380-630 region on a 1024 diode-intensified optical multichannel analyzer (Princeton Instruments, Inc., Princeton, NJ) with a resolution of 0.25 nm/diode. Each fluorescence emission spectrum was collected from 1 to 10 s. Data were stored and processed on an 80286 IBM PS/2 microcomputer using the Jobin-Yvon "Enhanced Prism" software. It should be noted that, in order to avoid any possible fluorescence from a plastic or glass support during analysis with near-UV excitation, cells were grown on quartz plates that were then placed on the microscope stage in 50-mm thermostated Petri dishes, filled with 5 ml of phosphate-buffered saline. A uranyl glass bar was used as a fluorescence standard to control laser power and instrumental response and to enable quantitative comparison between spectra recorded on different days. Sample heating, photobleaching, and photo damage were assessed empirically and found to be negligible under our experimental conditions. In particular, cells always remained viable after repeated fluorescence determinations, as controlled by phase-contrast micros.

### Iron affinity resin

Fe-NTA-Agarose was constructed from commercially available nickel NTA agarose (Qiagen, S.A., France). The Ni-NTA agarose was stripped with EDTA and recharged with iron as been described (64). Binding and pull down experiments were performed in 50 % ethanol and assumed 100% replacement of Ni sites with Fe^2+^ (as warranted by manufacturer). For binding experiments, the indicated amounts of curcumin were incubated with or without Fe-NTA-Agarose in 50% ethanol at room temperature (roughly 25 nmol metal binding sites in 0.5 ml total volume). After 10 minutes, the resin was removed by centrifugation, and aliquots of supernatants were diluted as needed (10 fold) to determine remaining curcumin in solution (absorbance at 435 nm). Affinity estimations were based on the method of Scatchard, assuming free curcumin equal to the amount remaining in solution in the presence of Fe-NTA-agarose, and bound curcumin equal to the amount curcumin added minus the free measured at each concentration shown. The experiments were conducted exactly as described by Messner et al. (52).

### Determination of lipid peroxidation and protein carbonylation

We used a lipid peroxidation assay kit (Abcam ab118970) to detect malondialdehyde (MDA) present in samples. The free MDA generated during lipid peroxidation refers to the oxidative degradation of lipids reacting with Thiobarbituric Acid (TBA) to generate a MDA-TBA adduct. The absorbance of MDA-TBA adduct was measured at 532 nm for a sensitivity as low as 1 nmol / well, and concentration calculated with provided standards. Protein carbonylation has been assayed using the Cayman’s Protein Carbonyl Fluorometric Assay Kit in Huh-7 cell lysates.

### Cell viability, mitochondrial membrane potential (ΔΨm), ROS and calcium levels

2 millions Huh-7cells were seeded on 6-well plates and maintained with 25 μM of curcumin for a given period of time ranging from 0 to 48h depending on the experiments. After treatment, cells were detached, harvested, washed and then resuspended together with their supernatants in Phosphate Buffer Saline (PBS). Before flow cytometry, various staining were used : DiOC_6_(3) was added at 40 nM final concentration for ΔΨ*m* determination, DCFH-DA at 5 μM for hydrogen peroxide detection, MitoSOX at 1 μM for superoxide anion detection. Most of the time a double staining is realized in order to simultaneously assess cell viability, with propidium iodide (PI; stock at 1mg.mL^−1^)in the case of DiOC_6_(3), DCFH-DA and Fluo4-AM and with 2 μg/ml TO-PRO-3 iodide (stock at 1mg.mL^−1^) for MitoSOX. A supplemental double staining was used for the distinction between viable, apoptotic and necrotic cells with YO-PRO-1 / PI in parallel with a Annexin-V / PI staining realized with the Annnexin-V-FITC when needed (Immunotech, Beckman-Coulter). All samples were measured and analyzed on a FACS Calibur 4C as previously described (73, 74).

### Detection of intracellular GSH activity and Glutathion peroxidase assay

Huh-7 cell content of GSH was determined using 50 μM monochlorobimane (mBCl, Molecular Probes, Eugene, OR) in an incubation of 20 min at room temperature in the dark (74). Monochlorobimane was dissolved in 100% ethanol to a stock concentration of 40 mM and stored at −20°C and special precautions were used to minimize the exposure to ambient light. To assess efficiency of mBCl for detection of intracellular GSH, Huh-7 were also incubated with N-ethylmaleimide (NEM, Sigma) as a control, a GSH depleting agent, which has been used previously to establish the specificity of mBCl for detection of GSH (75). N-ethylmaleimide was prepared as a stock solution in 100% ethanol and was added to cell suspension at 100 mM final concentration for 10 min at room temperature prior to addition of mBCl. Monochlorobimane fluorescence was assessed using a UV laser with excitation wavelength at 360 nm and an emission at 585 ± 42 nm (Fl-2) set on a FACS Aria (Becton-Dickinson, USA). For glutathion peroxidase (GPX4) activity determination we used a colorimetric assay from Abcam (ab102530) and follow manufacturer instructions. Data are expressed in enzymatic units per mg of protein.

### Calcein quenching experiments

In order to measure calcein-quenching, cells were loaded with 0.25 μg/ml calcein-1AM in serum free media for 30 minutes at 37°C, rinsed, and then treated (in triplicate) as specified in complete culture media for 2 hours. The cells were rinsed three times with PBS, and green fluorescence was measured by flow cytometry at 530 ± 30 nm.

### Caspase activation & fluorimetric assays

Isolated Huh-7 cells were washed and suspended in calcium-free buffer solution (140mM NaCl, 1.13mM MgCl_2_, 4.7mM KCl, 10mM glucose, 0.1 M EDTA, and 10mM HEPES, pH 7.2). Cells were then loaded at room temperature for 30 min with fluorescent indicator-linked substrates for activated caspase-8 (10 μM Z-IETD-R110; Molecular Probes), caspase-9 (10 μM Z-LEHD-R110; Molecular Probes), or caspases 3/7 (Caspase-3/7 Green ReadyProbes™ reagent with a DEVD sequence, Molecular Probes).

## ACKNOWLEDGEMENTS

As this manuscript is made with LATEX we thank the whole TEX community and the work of Dr. Ricardo Henriques’s lab for the initial available template.

## AUTHOR CONTRIBUTIONS

These contributions follow the Contributor Roles Taxonomy guidelines: https://casrai.org/credit/.

Conceptualization: N/A; Data curation: P.X.P; Formal analysis: P.X.P, N.E.R, A.M; Funding acquisition: P.X.P; Investigation: N.E.R, A.M, A.S, G.N, F.S, P.X.P; Methodology: A.M, P.X.P; Project administration: P.X.P; Resources: F.S, P.X.P; Software: N/A; Supervision: P.X.P; Validation: P.X.P; Visualization: N/A; Writing – original draft: N.E.R, A.M, P.X.P ; Writing – review & editing: All authors.

## COMPETING INTERESTS

The authors declare no competing interests.

